# Significant enrichment of Herpesvirus interactors in GWAS data suggests causal inferences for the association between Epstein Barr virus and multiple sclerosis

**DOI:** 10.1101/624049

**Authors:** Rosella Mechelli, Renato Umeton, Sundararajan Srinivasan, Arianna Fornasiero, Michela Ferraldeschi, IMSGC and WTCCC2, Diego Centonze, Cinthia Farina, Marco Salvetti, Giovanni Ristori

**Affiliations:** San Raffaele Roma Open University and IRCCS San Raffaele Pisana, Rome, Italy; Clinical & Translational Informatics Department, Dana-Farber Cancer Institute, Boston, USA; Institute of Experimental Neurology (INSpe) and Division of Neuroscience, IRCCS San Raffaele Scientific Institute, Milan, Italy; Centre for Experimental Neurological Therapies (CENTERS), Department of Neurosciences, Mental Health and Sensory Organs, Sapienza University of Rome, Italy; The International Multiple Sclerosis Consortium & the Wellcome Trust Case Control Consortium 2, Oxford, UK; IRCCS Istituto Neurologico Mediterraneo (INM) Neuromed, Pozzilli, Italy; Clinica Neurologica Dipartimento di Medicina dei Sistemi, University of Tor Vergata, Rome, Italy

**Keywords:** genes, Epstein-Barr virus, herpesvirus, interactome, multiple sclerosis, GWAS, gene expression, autoimmune disease

## Abstract

We exploited genetic information to assess the role of non-genetic factors in multifactorial diseases. To this aim we isolated candidate “interactomes” (i.e. groups of genes whose products are known to physically interact with environmental exposures and biological processes, plausibly relevant for disease pathogenesis) and analyzed nominal statistical evidence of association with genetic predisposition to multiple sclerosis (MS) and other inflammatory and non-inflammatory complex disorders. The interaction between genotype and Herpesviruses emerged as specific for MS, with Epstein Barr virus (EBV) showing higher levels of significance compared to Human Herpesvirus 8 (HHV8) and, more evidently, to cytomegalovirus (CMV). In accord with this result, when we classified the MS-associated genes contained in the interactomes into canonical pathways, the analysis converged towards biological functions of B cells, in particular the CD40 pathway. When we analyzed peripheral blood transcriptomes in persons with MS, we found a significant dysregulation of MS-associated genes belonging to the EBV interactome in primary progressive MS. This study indicates that the interaction between herpesviruses and predisposing genetic background is of causal significance in MS, and provides a mechanistic explanation for the long-recognized association between EBV and this condition.

## INTRODUCTION

Many studies on environmental exposures in multifactorial diseases are correlative and associative in nature, hence the need to focus on causality and mechanism (Fischbach, 2018). Genome-wide association studies (GWAS) have the capability to inform about the causal relevance of associations between environmental exposures and disease. However, the difficulty in extracting value from the study of gene-environment interactions limits the interpretation of GWAS, therefore hindering what could be a virtuous ‘genes-to-environment-to genes-again’ iterative learning process (Visscher et al., 2017).

In principle, Mendelian randomization allows to test for a causal effect of an epidemiological association. With this method, an environmental exposure is deemed to be plausibly causal if a genetic variant influencing the exposure is also directly associated with the disease (Davey Smith and Hemani, 2014). However, the influence of genetic variants on environmental exposures that associate with multifactorial diseases is far from being fully understood, particularly at the genome-wide level. More actionable data are available about the interaction, at the protein level, between human gene products and exposures. Furthermore, it has been shown that interacting disease-associated proteins have the propensity to influence each other in biologically relevant modules (Menche et al., 2015). Hence, in an approach akin to Mendelian randomization, we tested the possible causal significance of environmental exposures by measuring, at the genome-wide level, which genetic variants interacting with the exposure (i.e. influencing the exposure) were significantly enriched in MS GWAS data. At first, we began to explore this approach in an analysis limited to multiple sclerosis (MS) GWAS data, looking for statistical enrichments of associations among “candidate interactomes” (modules of genes whose products are known to physically interact with environmental factors of plausible, uncertain, or unlikely relevance for MS pathogenesis -Mechelli et al., 2013). This work provided a preliminary suggestion about the possible relevance of the interaction between host genotype and Epstein Barr virus (EBV) in MS.

Here we exploited the availability of new MS-GWAS data to replicate and refine the previous findings. Moreover, we extended the analysis to GWAS from other multifactorial diseases (type 1 and type 2 diabetes, rheumatoid arthritis, Crohn disease, celiac disease, bipolar disorder, hypertension, and coronary artery disease). Finally, we included updated and additional interactomes of diverse potential relevance for all the above conditions (see workflow in Figure1). These analyses confirmed the validity of our previous suggestion about herpesviruses as a component cause of MS and showed that this is not common to the other diseases considered in this study. A related issue is the clarification of the biological consequences of the observed associations. Elegant studies are resolving the functional implications of individual disease-associated variants (Gregory et al., 2012; Steri et al., 2017). Given the complex nature of multifactorial diseases (Wray et al., 2018), and particularly for translation into clinical benefit, a more general understanding of how gene-environment interactions generate gene expression alterations is also necessary (Gallagher and Chen-Plotkin, 2018). We therefore analyzed transcriptomes from the peripheral blood of persons with MS to verify whether genes that are nominally associated with MS, and are part of our interactomes, were more frequently present among dysregulated genes (see workflow in Figure1). Data showed that the expression of MS susceptibility genes, whose products interact with EBV, appeared to be more frequently dysregulated than expected in primary progressive MS. These results could be translated into meaningful biology as the analysis of the biological functions affected by MS-associated interactomes converged on the CD40 signaling, a main co-stimulatory pathway of B lymphocytes (the primary target of EBV infection) strongly implicated in MS pathophysiology (Smets et al., 2018).

**Figure 1:**
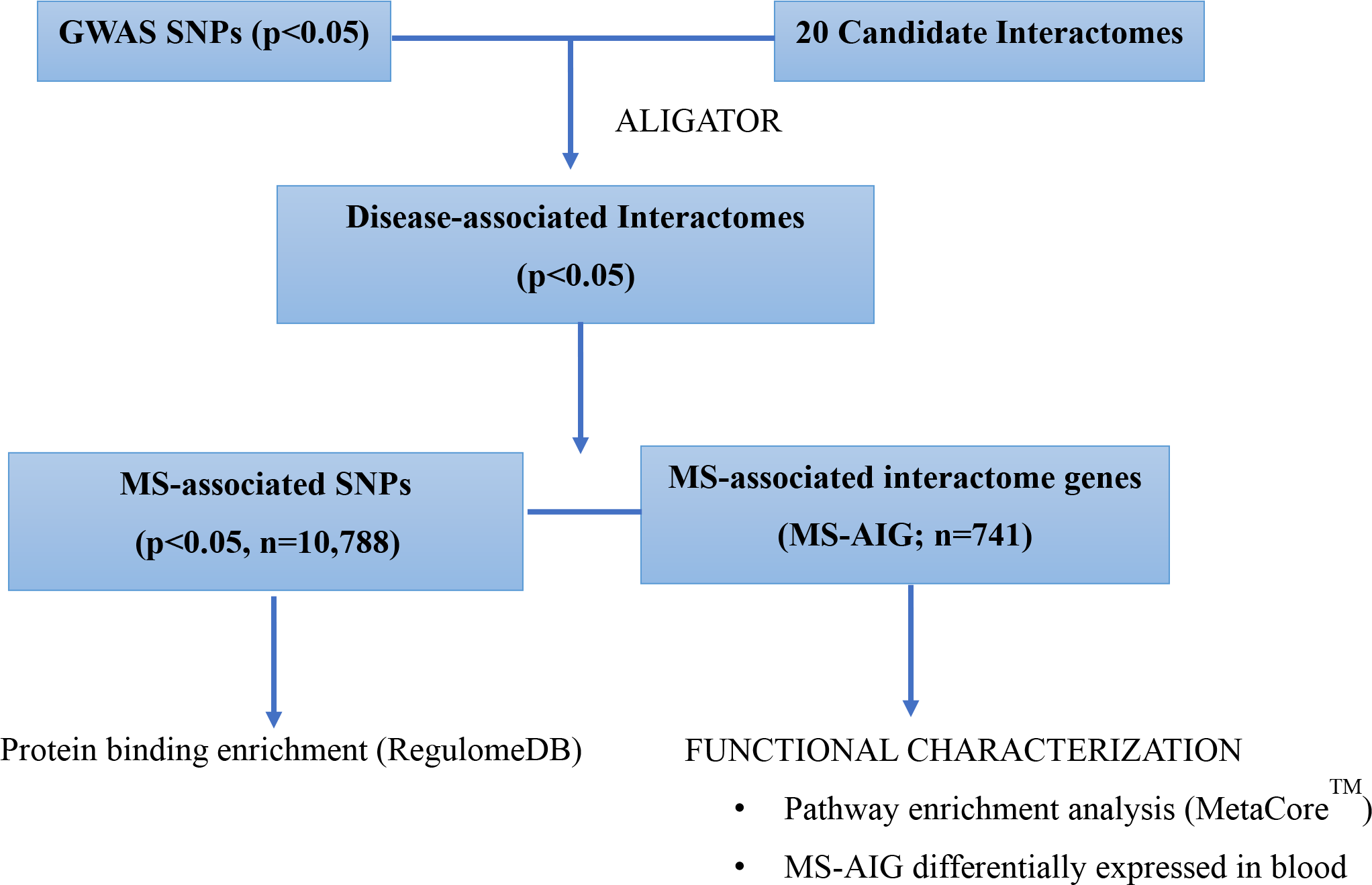
Schematic representation of the study.

## METHODS

The “candidate interactome” approach was applied as previously described (Mechelli et al., 2013). In brief, we obtained 20 candidate interactomes from the literature. Six were manually curated [Epstein-Barr virus (EBV), Hepatitis B virus (HBV), Cytomegalovirus (CMV), HHV8 Kaposi sarcoma virus (HHV8), JC virus (JCV), Inflammasome]; 14 were obtained from databases of molecular interactions and from published high-throughput experimental approaches: Autoimmune regulator (AIRE), Vitamin D receptor (VDR), Aryl hydrocarbon receptor (AHR), Sirtuin 1 (SIRT1), Sirtuin 7 (SIRT7) were obtained from BIOGRID (http://thebiogrid.org) (Chatr-Aryamontri et al., 2015), Polyomavirus was obtained from VirusMentha database (Calderone et al., 2015), Human-microRNA targets were obtained from miRecords database (Xiao et al., 2009), proteins targeted by 70 innate immune-modulating viral open reading frames from 30 viral species (VIRORF, Pichlmair et al., 2012), Human immunodeficiency virus (HIV, Jager et al., 2011), Hepatitis C virus (HCV, de Chassey et al., 2008), human innate immunity interactome for type I interferon (h-IFN, Li et al., 2011), H1N1-influenza (H1N1, Shapira et al., 2009), histone deacetylases (HDAC; Joshi et al., 2013), Chlamydia (Mirrashidi et al., 2015) (Table 1)).

**Table 1:** List of interactomes and related P-values of association with MS. In bold are indicated the statistically relevant associations.

| INTERACTOME | SIZE | SOURCE | P-VALUE<br>GWAS 2011 | P-VALUE<br>GWAS 2013 | P-VALUE<br>METACHIP1 | P-VALUE<br>METACHIP2 |
| --- | --- | --- | --- | --- | --- | --- |
| AHR | 38 | BioGRID | 0.5428 | **0.0260** | 0.3744 | 0.5332 |
| AIRE | 80 | BioGRID | **0.0342** | 0.2412 | 0.181 | 0.1276 |
| Chlamydia | 329 | Experimental data | 0.9910 | **0.0010** | 0.9864 | 0.982 |
| CMV | 96 | Manually curated | **0.0444** | 0.6720 | **0.0482** | **0.0298** |
| EBV | 261 | Manually curated | **0.0012** | 0.0828 | **0.0016** | **0.0004** |
| H1N1 | 87 | Experimental data | 0.9572 | 0.4228 | 0.9738 | 0.9516 |
| HBV | 123 | Manually curated | **0.0170** | 0.9636 | 0.1504 | 0.1668 |
| HCV | 202 | Experimental data | 0.4244 | 0.1758 | 0.3216 | 0.2818 |
| HDACs | 287 | Experimental data | 0.9610 | 0.2060 | 0.9562 | 0.9126 |
| HHV8 | 149 | Manually curated | **0.0046** | 0.2094 | **0.0052** | **0.0022** |
| HIV | 446 | Experimental data | **0.0026** | 0.3970 | **0.0216** | **0.0256** |
| huIFN | 113 | Experimental data | 0.2176 | 0.3104 | 0.3452 | 0.5364 |
| Human-miRNA targets | 294 | miRecord | 0.1514 | 0.5590 | 0.4754 | 0.3818 |
| Inflammasome | 194 | Manually curated | 0.1470 | 0.3410 | 0.0722 | 0.051 |
| JCV | 10 | Manually curated | 1.0000 | 1.0000 | 1.0000 | 1.0000 |
| Polyomavirus | 56 | VirusMentha | 0.2536 | **0.0418** | 0.1912 | 0.1996 |
| SIRT1 | 285 | BioGRID | 0.3032 | 0.0724 | 0.1374 | 0.1542 |
| SIRT7 | 658 | BioGRID | 0.5146 | **0.0400** | 0.3138 | 0.3144 |
| VDR | 92 | BioGRID | 0.3020 | 0.5102 | 0.2302 | 0.3212 |
| VIRORF | 579 | Experimental data | 0.0610 | **0.0152** | 0.089 | 0.0704 |

Abbreviations: AHR= Aryl hydrocarbon receptor; AIRE= autoimmune regulator; BioGRID= Biological General Repository for Interaction Datasets; CMV= Cytomegalovirus; EBV= Epstein-Barr virus; HBV= Hepatitis B virus; HCV= Hepatitis C virus; HDACs=Histone deacetylases; HHV8= Human Herpesvirus 8; HIV= Human Immunodeficiency virus; H1N1= Influenza A virus; hu-IFN= human innate immunity interactome for type I interferon; Human-miRNA targets= gene targets for human miRNA; Inflammasome= multiprotein complex responsible for activation of inflammatory processes and pyroptosis; JCV= JC virus; SIRT1= Sirtuin 1; SIRT7= Sirtuin 7; VIRORF= Virus Open Reading Frame; VDR= vitamin D receptor.

The manually curated interactomes were obtained by selecting only those interactions that were reported by two independent sources or were confirmed by the same source with distinct experimental approaches. In all cases, we considered only physical-direct interactions (Mechelli et al., 2013). All SNPs which did not pass quality checks in the GWAS studies were filtered out from the original data. As reference to gather gene and single nucleotide polymorphism (SNP) details from their HUGO Gene Nomenclature Committee (HGNC) Ids and rsids, a local copy of the Ensembl Human databases (version 75, databases core and variation, including SNPs coming from the 1000 Genome project) was employed; the annotation adopted for the whole analysis was GRCh37-p13, that includes the release 6 patches (Genome Reference Consortium: human assembly data - GRCh37.p13 - Genome Assembly (www.ncbi.nlm.nih.gov/assembly/GCF_000001405.25). We used ALIGATOR (Association LIst Go AnnoTatOR, Holmans et al., 2009; Houlston et al. 2010) program to evaluate how SNPs and single genes get summed to provide total contribution of candidate interactomes (Mechelli et al., 2013). The idea behind ALIGATOR’s strategy is to evaluate gene category significance by means of a bootstrapping approach: compare each interactome with the null hypothesis, that was built using random permutations of the data based on a non-parametric bootstrap analysis that uses the Gene Ontology database. Filtering criteria for the SNP selection included (i) a p-value significance taken at p-value cut-off (P-CUT) in the summary statistics; (ii) linkage disequilibrium (LD) filtering was applied on SNPs that have an r2<0.2 and those that are farther than 1000kb. The parameters used to configure ALIGATOR are those reported in its reference paper (Holmans et al., 2009): P-CUT was taken at 0.05, and the SNP-gene association parameter was maintained at its default value of 20kbp. For all disease-associated interactomes, SNPs were extracted after the ALIGATOR filtering and bootstrapping analysis steps. The idea behind this extraction was to pinpoint which SNPs were considered relevant by the ALIGATOR algorithm. Genes were mapped to SNPs by the ALIGATOR internal mapping step, for bp distance, p-value, and LD, using the default settings optimized in Holmans et al. (Holmans et al., 2009).

### Protein binding enrichment

After the ALIGATOR analysis, we used RegulomeDB database (Boyle et al., 2012; that is fed with live data from ENCODE and Roadmap Epigenomics) to look for proteins binding in regions containing “post-match” SNPs that displayed nominally significant associations with MS. When SNPs are bound by two or more proteins a Fisher’s exact test (2×2 table), controlling q-values for multiple-testing [false discovery rate (FDR), Benjamini-Hochberg], was computed to evaluate whether this co-occurrence is statistically relevant.

### Network and pathway analysis

MetaCore© (version 6.29 build 68613; GeneGO, Thomson Reuters, New York, N.Y) was employed to classify genes, carrying SNPs obtained from ALIGATOR analysis, in “canonical pathways” and evaluate their possible enrichment in specific categories. Canonical pathway maps represent a set of signaling and metabolic maps covering human biological processes in a comprehensive way. All maps are created by Thomson Reuters scientists by a high-quality manual curation process based on published peer-reviewed literature and validation experiments (Bolser et al., 2012).

### Blood transcriptome

Human peripheral blood mononuclear cell (PBMC) microarray datasets analyzed in this study were published in Srinivasan et al., 2017a; Srinivasan et al., 2017b) and deposited at EBI Array express database. The dataset was generated by Illumina Human Ref-8 v2 microarrays and contained PBMC transcriptomes of persons with MS (46 with clinically isolated syndromes, 52 with relapsing-remitting, 23 with primary-progressive, 21 with secondary-progressive disease) and 40 healthy controls.

### Frequency of interactome genes in MS transcriptomes

To verify whether genes obtained after ALIGATOR analysis, MS-associated interactome genes (MS-AIG), were enriched in differentially expressed genes (DEG) in MS, we measured their frequency in DEG list and in the global transcriptome. We verified whether the frequency of MS-AIG was significantly higher in the DEG list than the expected frequency of post-match genes in a random selection of transcripts from the database, by chi-square test with Yates’s correction using Graph Pad.

## RESULTS

### Selection of candidate interactomes and GWAS data

We considered 20 environmental exposures or biological processes of variable relevance for MS and other autoimmune diseases. We manually curated or extracted from published databases the interactomes of 9 viruses (EBV, HBV, CMV, HHV8, JCV, HIV, HCV, H1N1, Polyomavirus), 1 bacterium (Chlamydiae), 9 protein/process hubs (AIRE, VDR, AHR, VIRORF, h-IFN, HDAC, SIRT1, SIRT7, Inflammasome) and 1 human miRNA-mRNA network.

The interactomes manually curated (CMV, EBV, HBV, HHV8 and JCV) or obtained from publicly available databases (AHR, AIRE and VDR), that we have already selected in our previous work (Mechelli et al. 2013), were updated. We constructed an inflammasome interactome analyzing the literature and selecting all the physical-direct interactions among inflammasome proteins and other cellular proteins. The HDACs interactome was obtained from a unique source where different methods had been used to highlight protein-protein interactions among the 11 known HDAC enzymes and other cellular proteins (Joshi P et al. 2013). We considered also other deacetylase enzymes belonging to the sirtuin family (SIRT1 and SIRT7), whose interactomes were obtained from BioGRID database (Stark et al., 2011). We obtained the human miRNA-mRNA interactome from miRecords, an integrated resource for microRNA–target interactions, taking into consideration only experimentally verified miRNA human targets (Xiao et al., 2009). The Chlamydiae interactome was obtained from Mirrashidi et al., that used affinity purification and mass spectrometry approaches to highlight interactions among bacterial and cellular proteins (Mirrashidi et al.,2015). Finally, VirusMentha, a protein-protein interaction database specific for virus-host interactions (Calderone et al.,2015), was used to obtain Polyomavirus interactome.

The MS-GWAS data came from the first data set published in 2011 (International Multiple Sclerosis Genetics Consortium & Wellcome Trust Case Control Consortium,2 2011; MS-GWAS 2011) and Immunochip data, published in 2013 (International Multiple Sclerosis Genetics Consortium, 2013; MS-GWAS 2013), in which SNPs selected from immune-related loci were especially considered. To globally evaluate the contribution of all MS-associated SNPs, we constructed two combinations of the MS-GWAS 2011 and MS-GWAS 2013: in the first combination, METACHIP1, MS-GWAS 2011 data were considered in case of overlap (i.e., if both chips had a SNP in the same position, the MS-GWAS 2011 p-value was preferred); in the second combination, METACHIP2, MS-GWAS 2013 data were preferred in case of overlap (Suppl. Figure 1).

We also considered the published GWAS data from other immune-mediated or other complex disorders (type 1 and type 2 diabetes, rheumatoid arthritis, Crohn disease, celiac disease, bipolar disorder, hypertension, coronary artery disease and related healthy controls) from Wellcome Trust Case Control Consortium,2 (WTCCC2). These data were re-worked in the context of our candidate interactome approach to verify the specificity of the findings for MS.

### Candidate interactome analysis

We selected significant SNPs associated with MS and other complex diseases (type 1 and type 2 diabetes, rheumatoid arthritis, Crohn disease, celiac disease, bipolar disorder, hypertension, coronary artery disease), from GWAS datasets, considering a p-value cut-off of association with each disease at 0.05.

We investigated their statistical enrichment of association within each one of the 20 selected interactomes using ALIGATOR (see workflow in Figure 1). Spearman’s coefficients were calculated to evaluate linear correlation between the size of single interactomes and their cumulative p-value of association with the diseases, but no correlation was found in any case, indicating that the size of interactomes does not influence discovery (Suppl. Figure 2-3). When analyzing the frequency of disease-associated genes from distinct MS GWAS studies in the 20 interactomes, viral interactomes were more reproducibly associated with MS (Table 1). The three herpes viruses studied showed statistical significance with good consistency throughout the analyses (levels of significance: EBV>HHV8>CMV), except for MS GWAS 2013. This result characterized MS with respect to all other conditions, including the immune-mediated disorders like diabetes, rheumatoid arthritis, Crohn disease and celiac disease (Table 2). In fact, significant associations were scattered in non-MS conditions and herpesvirus interactomes reached significance only in coronary artery disease (CMV) and type 2 diabetes (HHV8). Among other, non-herpes, viral interactomes, the one of HIV was MS-associated in the GWAS 2011, METACHIP1 and METACHP2 but not in GWAS 2013. Other viral interactomes showing associations with MS were HBV (GWAS 2011), polyomavirus (GWAS 2013) and VIRORF (GWAS 2013). Notably, also for these non-herpes, viral interactomes, the association with the other diseases was either absent (HBV and HIV) or present only in one condition (VIRORF in T2 diabetes).

**Table 2:** List of interactomes and related P-values of association with other diseases. In bold are indicated the statistically relevant associations.

| INTERACTOME | SIZE | SOURCE | CHRON DISEASE | RHEUMATOID ARTHRITIS | CELIAC DISEASE | T1 DIABETES | T2 DIABETES | HYPERTENSION | BIPOLAR DISORDER | CORONARY ARTERY DISEASE |
| --- | --- | --- | --- | --- | --- | --- | --- | --- | --- | --- |
| AHR | 38 | BioGRID | 0.8734 | 0.9176 | 0.3814 | 0.8262 | 0.3552 | 0.8256 | 0.892 | 0.8538 |
| AIRE | 80 | BioGRID | 0.711 | 0.274 | 0.658 | 0.7804 | 0.9188 | **0.0072** | 0.5934 | 0.99 |
| Chlamydia | 329 | Experimental data | 0.9266 | 0.9818 | 0.7758 | 0.1918 | 0.9464 | 0.6354 | 0.8938 | 0.201 |
| CMV | 96 | Manually curated | 0.2198 | 0.5126 | 0.0704 | 0.9056 | 0.3438 | 0.6956 | 0.5378 | **0.0288** |
| EBV | 261 | Manually curated | 0.4578 | 0.523 | 0.4716 | 0.9968 | 0.5444 | 0.1674 | 0.539 | 0.4784 |
| H1N1 | 87 | Experimental data | 0.7792 | 0.6486 | 0.9022 | 0.5498 | 0.1316 | 0.688 | 0.693 | 0.7376 |
| HBV | 123 | Manually curated | 0.6862 | 0.7642 | 0.3258 | 0.5804 | 0.5576 | 0.8778 | 0.3766 | 0.7486 |
| HCV | 202 | Experimental data | 0.7698 | 0.5986 | 0.7654 | 0.8968 | 0.5666 | 0.9428 | 0.8824 | 0.9596 |
| HDACs | 287 | Experimental data | 0.886 | 0.6372 | 0.5472 | 0.772 | 0.9158 | 0.3306 | 0.368 | 0.9486 |
| HHV8 | 149 | Manually curated | 0.8732 | 0.6424 | 0.5574 | 0.3588 | **0.0214** | 0.7842 | 0.6578 | 0.3154 |
| HIV | 446 | Experimental data | 0.7636 | 0.903 | 0.6046 | 0.4518 | 0.0832 | 0.5958 | 0.7344 | 0.7496 |
| huIFN | 113 | Experimental data | 0.616 | 0.968 | 0.673 | 0.1132 | 0.7168 | 0.177 | 0.2146 | 0.6716 |
| Human-miRNA targets | 294 | miRecord | 0.0506 | 0.3044 | **0.0036** | 0.7706 | 0.8784 | 0.9244 | 0.9534 | 0.2258 |
| Inflammasome | 194 | Manually curated | 0.7568 | 0.093 | 0.187 | 0.3338 | 0.1026 | 0.3308 | 0.5632 | 0.7648 |
| JCV | 10 | Manually curated | 0.4108 | 1 | 0.1758 | 0.8256 | 0.1758 | 0.3734 | 0.836 | 0.0912 |
| Polyomavirus | 56 | VirusMentha | 0.8052 | 0.8536 | 0.1856 | 0.7474 | 0.1752 | 0.1268 | 0.9238 | 0.2774 |
| SIRT1 | 285 | BioGRID | 0.1684 | 0.7826 | 0.373 | 0.7812 | 0.727 | 0.9478 | 0.6784 | 0.0538 |
| SIRT7 | 658 | BioGRID | 0.1894 | 0.981 | 0.4086 | 0.936 | 0.1748 | 0.4836 | 0.1136 | 0.9566 |
| VDR | 92 | BioGRID | 0.9458 | 0.9886 | 0.1324 | 0.953 | 0.8164 | 0.4766 | 0.7634 | 0.8654 |
| VIRORF | 579 | Experimental data | 0.1976 | 0.7448 | 0.4766 | 0.8866 | **0.0372** | 0.7168 | 0.3362 | 0.2036 |

Abbreviations: AHR= Aryl hydrocarbon receptor; AIRE= autoimmune regulator; BioGRID= Biological General Repository for Interaction Datasets; CMV= Cytomegalovirus; EBV= Epstein-Barr virus; HBV= Hepatitis B virus; HCV= Hepatitis C virus; HDACs=Histone deacetylases; HHV8= Human Herpesvirus 8; HIV= Human Immunodeficiency virus; H1N1= Influenza A virus; hu-IFN= human innate immunity interactome for type I interferon; Human-miRNA targets= gene targets for human miRNA; Inflammasome= multiprotein complex responsible for activation of inflammatory processes and pyroptosis; JCV= JC virus; SIRT1= Sirtuin 1; SIRT7= Sirtuin 7; VIRORF= Virus Open Reading Frame; VDR= vitamin D receptor.

### Regulatory elements and pathways common to MS-associated interactomes

To better understand the functional picture that may emerge from the above results we evaluated the possibility that common regulatory mechanisms exist among all the MS-associated interactomes. To verify this hypothesis, we considered the SNPs and the genes resulting from the ALIGATOR analysis, that were responsible for the statistical enrichments of association of interactomes with MS: n=10,788 nominally significant SNPs and the corresponding 741 MS-AIG. We then used RegulomeDB database (Boyle et al., 2012) to look for proteins whose binding close to SNPs extracted from ALIGATOR analysis is enriched. We overlapped the SNPs with the protein binding map generated by ENCODE and RoadMap Epigenomics (Boyle et al., 2012), obtaining a list of enriched proteins. RNA Polymerase II Subunit A (POLR2A) and CCCTC-Binding Factor (CTCF) displayed the highest regulatory potential (Figure 2). Since each SNP can be bound by multiple proteins, we carried out an analysis of “co-occurrence” for POLR2A and CTCF evaluated using Fisher’s exact test. After controlling P-values for multiple-testing (FDR, Benjamini-Hochberg), co-occurrence of POLR2A and CTCF was significant (q<0.05).

**Figure 2:**
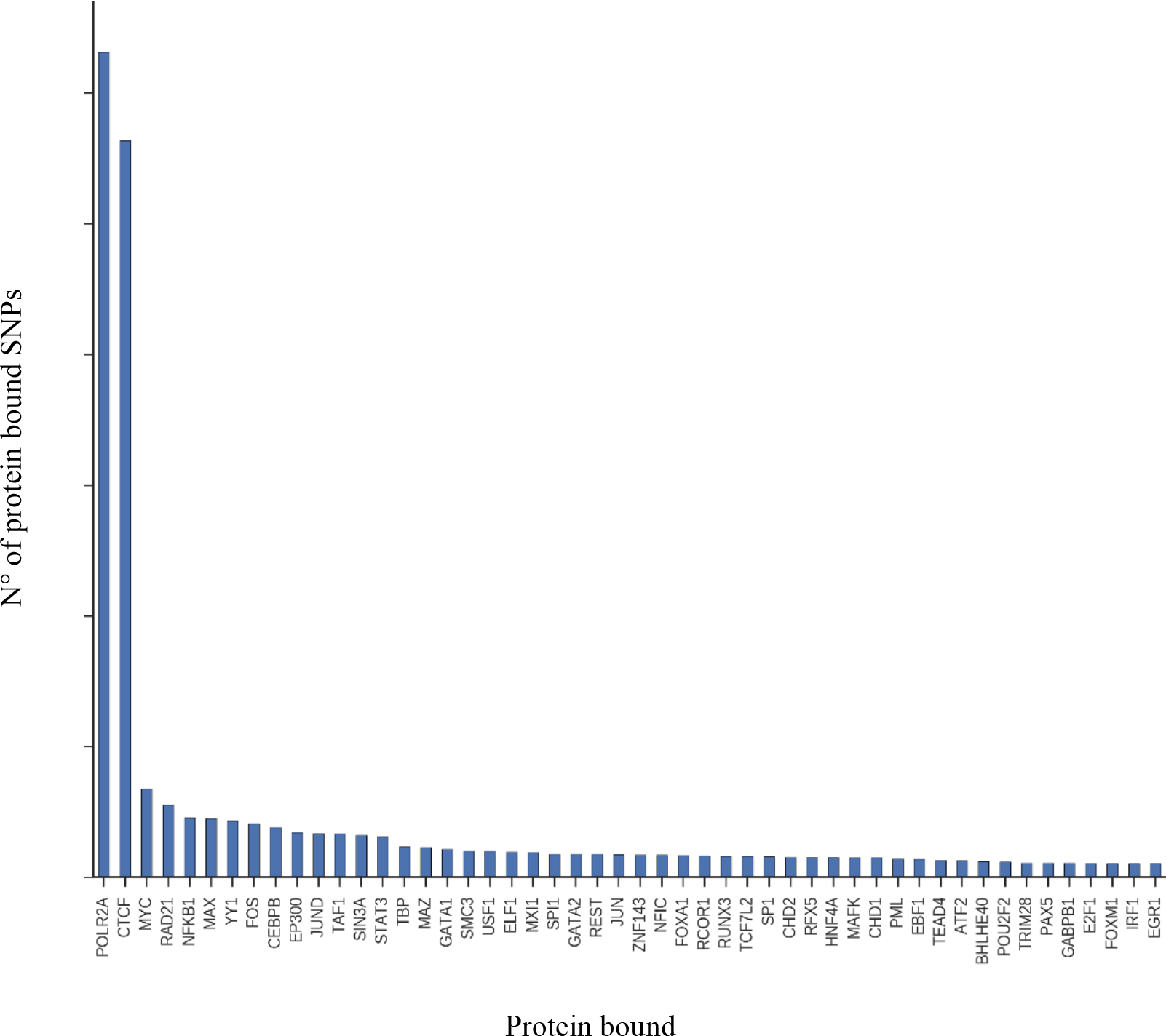
Protein binding enriched near MS-associated SNPs. The box plot displays the number of SNPs bound by a given protein (Y-axis) and the top 50 proteins enriched on MS-associated SNPs (x-axis).

We used MetaCore© to highlight the potential biological functions affected by the MS-AIG. The 741 MS-AIG were mapped to the internal MetaCore KnowledgeBase and a classification in canonical pathways was obtained. This analysis showed that the most critical biological process involved in MS through gene-environment interactions was the CD40 signaling, followed by stress-induced antiviral cell response, oxidative stress and apoptosis (Figure 3).

**Figure 3:**
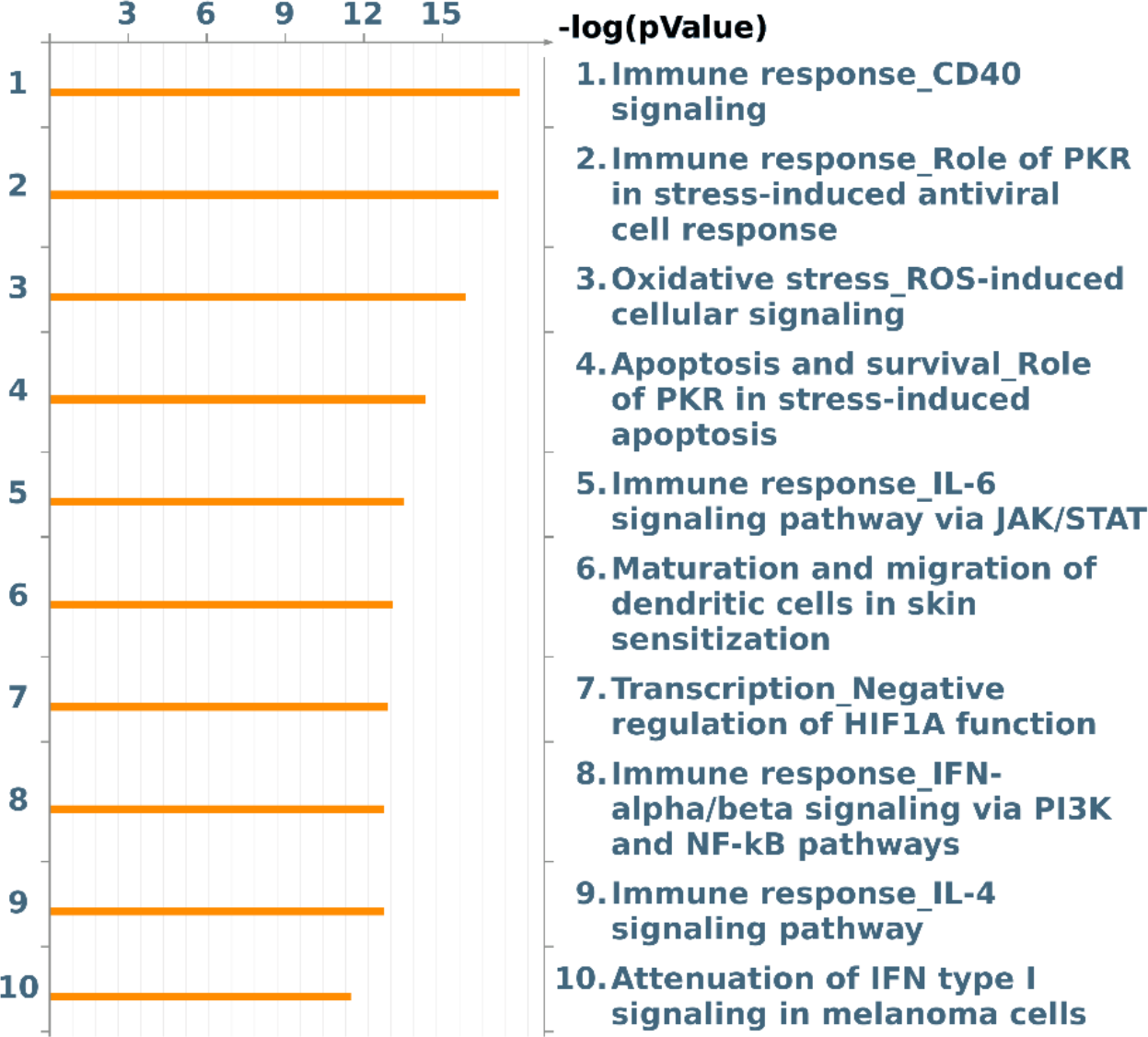
Genomic distribution and pathway enrichment analysis of MS-AIG. Top 10 pathways obtained from the analysis of the MS-AIG. The pathway analysis was performed using MetaCore and its canonical pathway maps. The lower p-value (-log Pvalue) means higher relevance of the MS-AIG within the pathway datasets.

### MS-AIG differentially expressed in blood transcriptome

To investigate the functional consequences of these findings we matched the 741 MS-AIG with gene expression data obtained from the peripheral blood of healthy subjects and persons with different MS phenotypes, including clinically isolated syndrome (CIS), relapsing-remitting (RR), secondary progressive (SP) and primary progressive (PP) MS (Srinivasan et al., 2017b). We calculated whether the MS-AIG were significantly enriched in the lists of genes differentially expressed in each MS forms. We took into consideration differentially expressed genes with p<0.05 and fold change >1.5. Totally, 110 MS-AIG were differentially expressed in at least one MS stage but no significant enrichments appeared in any of the 4 disease subtypes (Table 3). However, when considering separately the MS-AIG from single environmental interactomes, we observed an enrichment of differentially expressed MS-AIG from AIRE interactome in CIS patients, from AIRE and SIRT7 interactomes in RRMS and from EBV interactome in PPMS (Table 4).

**Table 3:** MS-AIG differentially expressed in blood. Frequency of MS-AIG in DEG in peripheral blood in distinct MS phenotypes.

| MS phenotypes | Interactome DE Probes (p<0.05) | MS-AIG matching blood transcript | Frequency of interactomeDE probes (%) | Global DEG (P<0.05) | Expressed probes | Frequency of global DEG (%) | Chi square | P-value |
| --- | --- | --- | --- | --- | --- | --- | --- | --- |
| CIS | 166 | 674 | 24.6 | 2636 | 10314 | 25.56 | 0.28 | 0.59 |
| RRMS | 170 |  | 25.2 | 2440 |  | 23.66 | 0.89 | 0.34 |
| PPMS | 249 |  | 36.94 | 3489 |  | 33.8 | 3.019 | 0.082 |
| SPMS | 226 |  | 33.56 | 3340 |  | 32.3 | 0.37 | 0.53 |

CIS = clinically isolated syndrome; RR-MS = Relapsing-Remitting multiple sclerosis; PP-MS = Primary progressive multiple sclerosis; SP-MS = Secondary progressive multiple sclerosis; DEG = differentially expressed genes.

**Table 4:** Frequency of interactome-specific MS-AIG in blood DEG. The enrichment of MS-AIG was calculated considering separately the genes obtained from each MS-associated interactome. The analysis was performed in distinct MS phenotypes. The associations reaching significance are highlighted in bold.

| Frequency of interactome DEG (%) |  |  |  |  |  |  |  |  |  |  |  |  |
| --- | --- | --- | --- | --- | --- | --- | --- | --- | --- | --- | --- | --- |
| Interactome | AHR | AIRE | CHLAMYDIA | CMV | EBV | HBV | HHV8 | HIV | POLYOMAVIRUS | SIRT7 | VIRORF | GLOBAL<br>TRANSCRIPTOME |
| MS<br>phenotypes |  |  |  |  |  |  |  |  |  |  |  |  |
| CIS | 25.00 | **50** | 29.03 | 34.62 | 24.32 | 16.00 | 22.09 | 21.43 | 45.45 | 27.93 | 30.51 | 25.56 |
| RRMS | 25.00 | **40.63** | 25.81 | 19.23 | 22.97 | 24.00 | 27.27 | 21.90 | 45.45 | **34.23** | 27.97 | 23.66 |
| PPMS | 25.00 | 40.63 | 38.71 | 38.46 | **44.59** | 46.00 | 42.86 | 32.86 | 27.27 | 31.53 | 34.75 | 33.83 |
| SPMS | 31.25 | 37.50 | 27.42 | 32.69 | 33.11 | 28.00 | 37.66 | 32.86 | 9.09 | 30.63 | 34.75 | 32.38 |

CIS = clinically isolated syndrome; RR-MS = Relapsing-Remitting multiple sclerosis; PP-MS = Primary progressive multiple sclerosis; SP-MS = Secondary progressive multiple sclerosis; DEG = differentially expressed genes.

## DISCUSSION

Our results strengthen the causal inference of many previous sero-epidemiological investigations showing associations between viruses and MS (Marrodan e al. 2019). In fact, we not only replicate the previous result (Mechelli et al., 2013), but does so in an increasingly meaningful setting that takes into account new genetic data. Also, the analysis of new “candidate interactomes”, that were not part of the exploratory study, reinforces this result. The surfacing of other herpesviruses, and the turn down of other viruses (e.g. HBV) that were significant in the previous study, are also in line with the sero-epidemiological literature, with obvious biological similarities between EBV and other herpesviruses, and – to some extent – with biological plausibility. Furthermore, the results indicate that this kind of analyses become increasingly meaningful and reliable as new genetic and protein-protein interaction data become available. It is remarkable that, in a study focused on the enrichment of susceptibility genes for psychiatric and neurological diseases in the Herpes simplex virus 1 interactome, (Carter, 2013) MS emerged as the disorder with the highest enrichment. Though this study was focused on HSV-1, and examined association of genes in the range of GWAS significance (and not p values of all SNPs with nominal significance as in our case), EBV emerged as the most significant pathogen-related pathway in MS. Furthermore, of the six recently identified genes harboring MS-associated low-frequency coding variants (International Multiple Sclerosis Genetics Consortium, 2018), five are either EBV interactors (PRKRA, TYK2, Geiger et al., 2006) or implicated in EBV-related pathophysiologies (PRF1 mutations are causative for EBV-associated lymphoproliferative disorders and hemophagocytic lymphohistiocytosis, Ono et al., 2018; HDAC7 is involved, more than other histone deacetylases, in the control of signal transduction and transcription factor modification during reactivation of EBV from latency, Bryant et al., 2002; GALC has tumor-suppressive effects in EBV-associated nasopharyngeal carcinoma, Zhao et al., 2015). For the sixth, NLRP8, there is no interaction reported in BIOGRID. Finally, the SNP with the highest association with white matter lesion volume, in the recent genome-wide association study of brain imaging phenotypes of the UK Biobank (Elliott et al., 2018), is in the 5’ UTR of EFEMP1, a gene that codes for another EBV interactor (its product interacts with two EBV proteins, BFLF2 and BRRF1; Gulbahce et al., 2012). This convergence of results reinforces the idea that this method of analysis can clarify the sense of observed “environmental” associations in the light of genetic information (indeed, performing these analyses on multiple genetic variants, which are likely to have a mechanistically direct effect on the exposure, increases the reliability of this kind of approaches; O’Connor and Price, 2018). Notably, when we looked at the biological functions that may be affected by the 741 MS-AIG, the CD40 signaling prevailed. CD40 is a main co-stimulatory pathway in B lymphocytes (the primary target of EBV infection), recently described as altered by MS risk variants (Smets et al., 2018), with possible consequences on the creation of memory B cells (Baker et al., 2018) and the activation of autoreactive CD4+ T lymphocytes (Jelcic et al., 2018; Baker et al., 2018). Our bioinformatics reworking on GWAS data disclosed an unanticipated relationship between CD40 and plasminogen activator gene in MS pathophysiology (La Starza et al. 2019).

Besides MS, in this study we applied the same analysis to other immune-mediated diseases where epidemiological studies have found associations with EBV, though probably less robust compared to MS. In these conditions, our data do not support the idea that the interaction between EBV and the genetic background of the host may be decisive for disease etiology. Some differences in the response to therapeutic interventions between MS and other immune-mediated diseases may be, at least in part, in accord with differences in the contribution of EBV (or other viruses) to disease pathophysiology. In particular, MS and RA greatly differ in their clinical response to tumor necrosis factor (TNF) antagonists, which worsen the former (in some cases even induce an MS-like demyelination in patients treated for other diseases) while being a mainstay for the treatment of the latter (and beneficial in other immune-mediated conditions including Crohn’s disease, ankylosing spondylitis and psoriasis). Among other possible explanations, including those linked to peculiarities of the TNF system in the inflamed brain, blocking or reducing the effects of TNF (one of the most ancient antiviral cytokines) may have detrimental effects in a disease that has a viral component. This may be particularly true in subjects harboring a genetic variant of the TNF-receptor 1 that mimics a TNF antagonist and is associated with MS but not with RA, psoriasis and Crohn disease. Conversely, immunotherapies that exploit cytokines with antiviral actions, such as interferon-beta, are effective in MS but not in RA (van Holten et al., 2005; Genovese et al., 2004) or in other immune-mediated diseases where they may concur in the development of the diseases (Axtell et al., 2011). Also, the response to anti-B lymphocyte therapies in MS, including in a progressive form of the disease, may be indicative of a role of EBV in disease pathophysiology as this virus remains for the lifetime of the host in memory B cells. Of note, in RA, anti-B lymphocyte therapies may not be as effective as anti-TNF treatments (Gonzalez-Vacarezza et al., 2014). In spite of these differences, other mechanisms of EBV involvement in autoimmunity cannot be excluded. For example, the binding of EBNA2 to genomic intervals associated with various autoimmune diseases, including MS, suggests another mechanism that operates across diseases (Harley et al., 2018; Ricigliano et al., 2015). Furthermore, also in view of recent results about HHV-6 and HHV-7 in Alzheimer’s disease and in a non-human primate model of MS (Readhead et al., 2018; Leibovitch et al. 2018) it will be necessary to study more immune-mediated diseases and various other “candidate interactomes” of viral origin to draw a complete picture of the long-suspected role of viruses in autoimmunity and in neurodegeneration.

The match of MS-AIG with gene expression lists from the peripheral blood of patients emphasizes AIRE interactors in CIS and RRMS, a not unexpected finding, considering the AIRE function in the maintaining of tolerance to tissue (auto)-antigens (Anderson et al. 2016; Zhu et al. 2016). Further studies will be needed to clarify the impact of SIRT7 interactors on RR-MS, especially in light of its involvement in inflammatory signaling pathways (Mendes et al 2017). The significant association of the EBV interactome in PP-MS confirms, at least in part, the results obtained by ALIGATOR analysis (see Table 1). To refine this finding, new studies will have to investigate the relationship between gene expression dysregulation and genetic background, not only of the host, but also of the viral agent as some EBV genomic variants appear to be associated with MS more than others (Mechelli et al., 2015). Moreover, it will also be important to understand how POLR2A and CTCF, that we identified as common regulatory mechanisms upstream the disease-associated interactomes, coordinate with all other transcription factors that have been mapped on a substantial fraction of autoimmune disease loci (Harley et al., 2018). Other groups observed the binding of CTCF in some specific genomic regions, bearing disease-associated SNPs (Zuvich et al., 2011; Martin et al., 2011). Our analysis extends this association to all SNPs that we found to be MS-associated in the interactome analysis. This is of interest considering data showing that CTCF frequently interacts with sites adjacent to Epstein-Barr nuclear antigen 2 (EBNA2) super-enhancer (Zhou H et al. 2015). Moreover, POLR2A and CTCF have recently been associated with risk SNPs in schizophrenia (Huo et al. 2019), suggesting a wider relevance of these two proteins for CNS diseases. Our study indicates that the interaction between host genotype and Herpesviruses (including EBV) is relevant for multiple sclerosis etiology, reinforcing the rationale for experimental medicine approaches aimed at advancing therapies while proving the causality of viral agents in MS. The results of a recent study on EBV-specific adoptive T cell therapy in patients with progressive MS support this perspective (Pender et al. 2018).

## Acknowledgments

This work was supported by Italian Foundation of Multiple Sclerosis (grant number 2014/R/12 to RM).

## Author contributions

R.M., R.U., G.R., M.S. conceived and designed the experiments. R.M. selected and constructed the candidate interactomes. R.U. performed the bioinformatics works. R.M., R.U., A.F., M.F., D.C., G.R., M.S. analyzed the data. S.S. and C.F. provided blood transcritptome data and were involved in data analysis. R.M., G.R., M.S. provided data interpretation and wrote the manuscript. C.F. and D.C. provided critical reading of the manuscript. IMSGC and WTTTC2 provided GWAS data.

## Competing interests

Mechelli R., Renato U., Fornasiero A., Ferraldeschi M., Ristori G. and Srinivasan S. declare no conflict of interest. Centonze D. is an Advisory Board member of Almirall, Bayer Schering, Biogen, GW Pharmaceuticals, Merck Serono, Novartis, Roche, Sanofi-Genzyme, Teva and received honoraria for speaking or consultation fees from Almirall, Bayer Schering, Biogen, GW Pharmaceuticals, Merck Serono, Novartis, Roche, Sanofi-Genzyme, Teva. He is also the principal investigator in clinical trials for Bayer Schering, Biogen, Merck Serono, Mitsubishi, Novartis, Roche, Sanofi-Genzyme, Teva. His preclinical and clinical research was supported by grants from Bayer Schering, Biogen Idec, Celgene, Merck Serono, Novartis, Roche, Sanofi-Genzyme e Teva

Farina C. received research support from Merck-Serono, Teva, Novartis.

Salvetti M. received research support and consulting fees from Biogen, Merck-Serono, Novartis, Roche, Sanofi, Teva

