## Supplementary figures 1-2-3 for "Significant enrichment of Herpesvirus interactors in GWAS data suggests causal inferences for the association between Epstein Barr virus and multiple sclerosis"

**Suppl. Figure 1**: **Schematic representation of METACHIP datasets construction**

Metachip 1 and 2 were built as the position-wise union of GWAS and Immunochip. Each dataset is represented by a line in wich sample SNPs are indicated with different shapes and colors depending on their source dataset (black round for SNPs coming from GWAS and empty round for SNPs coming from Immunochip). When a given SNP existed in both GWAS and Immunochip, we had to choose to which of the two we would give preference: Metachip1 gives preference to GWAS, while Metachip2 gives preference to Immunohip.

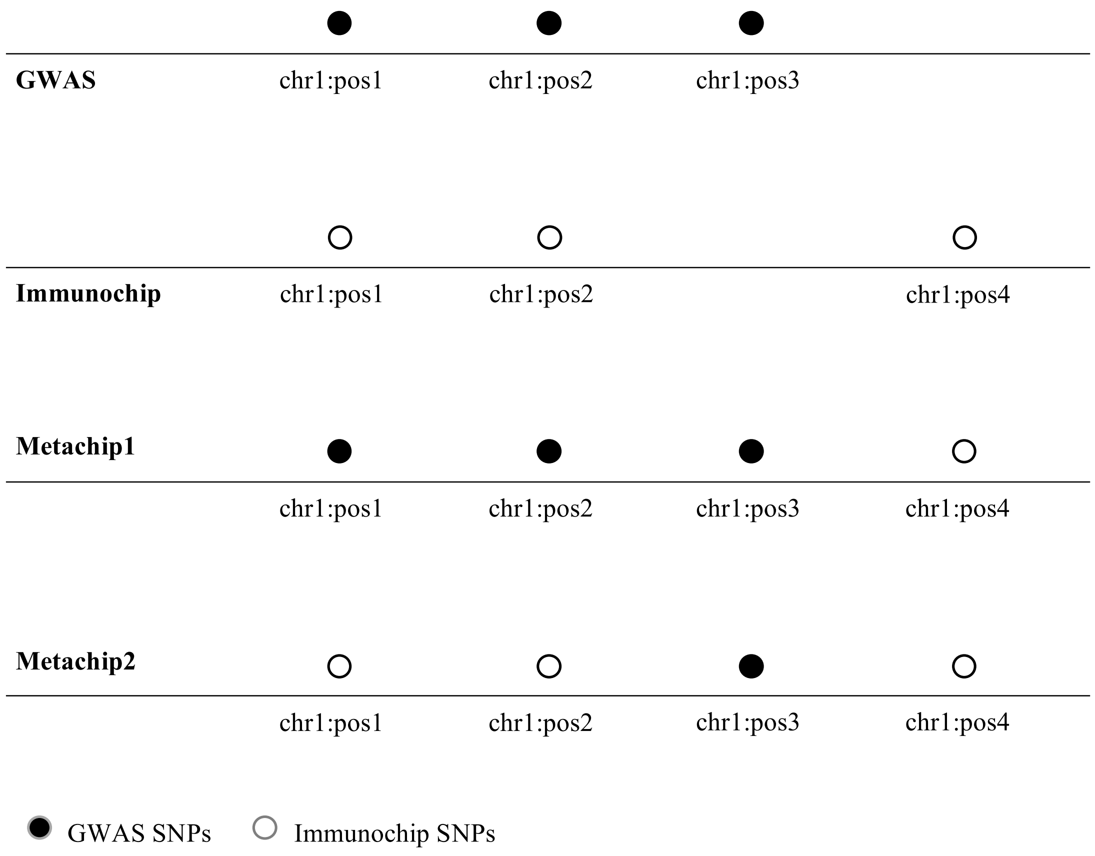

**Suppl. Figure 2: Spearman correlation in MS GWAS**

Linear correlations between the size of single interactomes and their cumulative p-value of association with MS calculated for each GWAS dataset (GWAS 2011, GWAS 2013, METACHIP 1 and METACHIP 2). The 95% confidence intervals of the Spearman correlation are greyed out and in all the analysis.

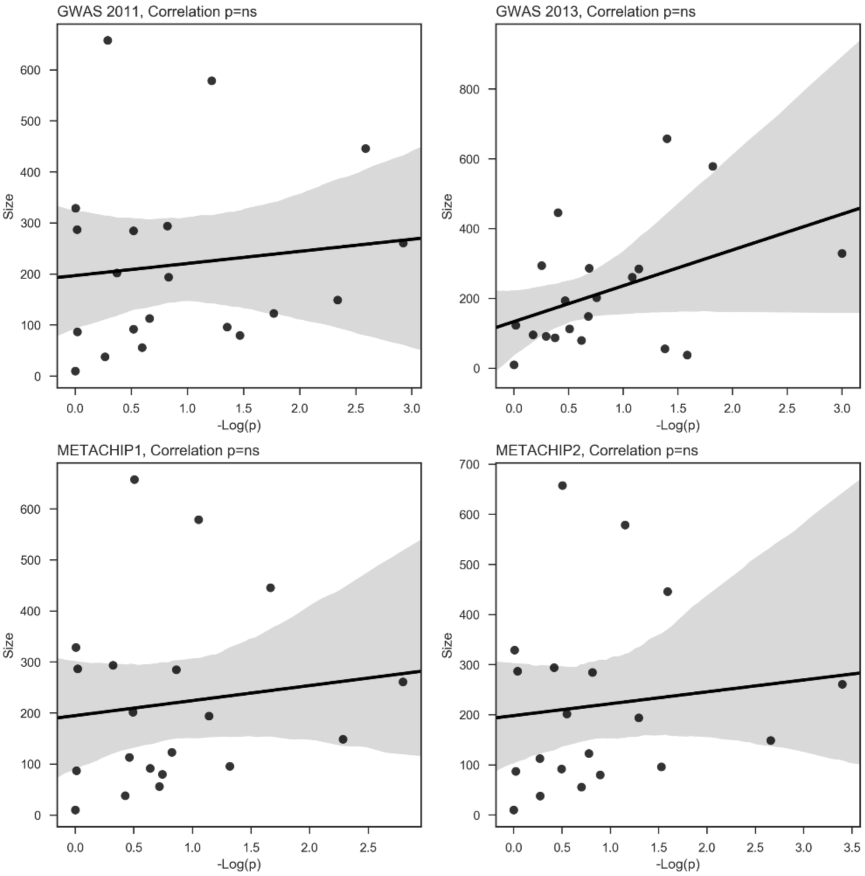

**Suppl. figure 3: Spearman correlation in non-MS GWAS**

Linear correlations between the size of single interactomes and their cumulative p-value of association with complex diseases calculated for each GWAS dataset (Bipolar disorder, celiac disease, Chron disease, coronary artery disease, hypertension, rheumatoid arthritis, T1 diabetes, T2 diabetes). The 95% confidence intervals of the Spearman correlation are greyed out and in all the analysis.

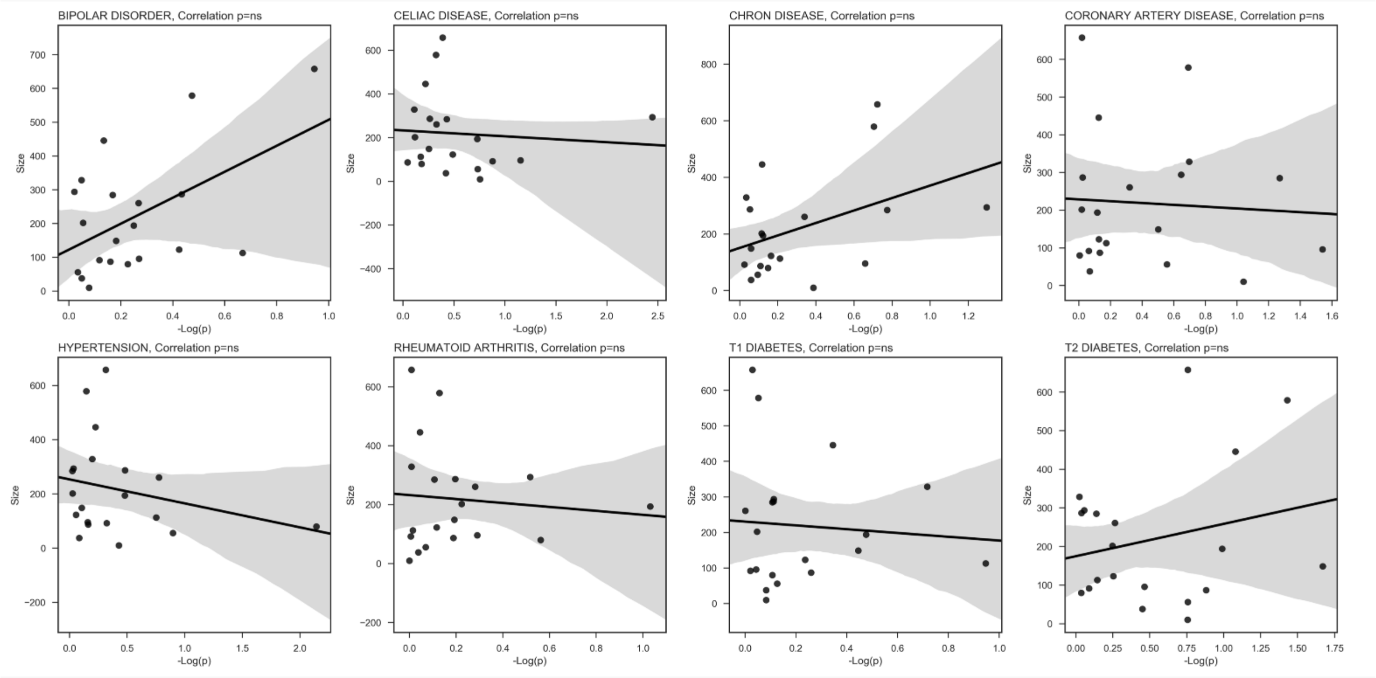
